# The *Drosophila* myogenic inhibitor *Him* gene is essential for adult muscle function and muscle stem cell maintenance

**DOI:** 10.1101/2024.09.06.611611

**Authors:** Robert Mitchell-Gee, Robert Hoff, Kumar Vishal, Daniel Hancock, Sam McKitrick, Cristina Newnes-Querejeta, TyAnna L. Lovato, Richard M. Cripps, Michael V. Taylor

**Author notes:** These authors contributed equally (Robert Mitchell-Gee, Robert Hoff).

## Abstract

Vertebrate muscle fibres have a population of Muscle Stem Cells (MuSCs), or “satellite cells”, vital to muscle growth, homeostasis and repair. In *Drosophila*, adult MuSCs with similar characteristics have only recently been described. This has opened up the *Drosophila* system for analysing how MuSCs operate in muscle maintenance, repair and ageing. Here we show that the *Him* gene is expressed in the adult muscle progenitors (AMPs), or myoblasts, that make the adult *Drosophila* thoracic flight and jump muscles. Notably, we also show that Him is expressed in the flight muscle MuSCs identifying *Him* as only the second genetic marker of these insect MuSCs. We then explored *Him* function. *Him* mutants have disrupted organisation of the thoracic jump muscle, resulting in reduced jumping ability. *Him* mutants also have a reduced pool of the myoblasts that will develop into the flight muscles. In the flight muscles themselves, *Him* mutants have an age-dependent decrease in the number of MuSCs, indicating that *Him* is required for maintenance of the adult muscle stem cell population. Moreover, this decrease in MuSCs coincides with a functional effect: there is an age-dependent decline in flight ability. Overall, *Him* is a novel marker of the *Drosophila* adult MuSC, and is required during ageing both to maintain MuSC number and flight ability.

## Introduction

The fruit fly *Drosophila melanogaster* has proven a valuable model for researchers to explore the genetic and cellular basis of muscle development (Dobi *et al*. 2015; Laurichesse and Soler 2020). During development, *Drosophila* undergoes two waves of skeletal muscle myogenesis; the first during embryogenesis gives rise to the larval musculature that is used until pupation; the second wave, during pupation, forms the diverse array of muscles found in the adult fly that function for two to three months. The different adult muscles arise from adult muscle progenitor cells (AMPs), a stem cell population that is put aside during embryogenesis and which then proliferates during larval life. The adult muscles include the thoracic indirect flight muscles (IFMs), which develop from wing disc-associated AMPs, and the jump muscle (also known as TDT, tergal depressor of trochanter), which develops from AMPs associated with the T2 mesothoracic leg disc (Jaramillo *et al*. 2009).

In recent years, muscle progenitor cells have been discovered in *Drosophila* at the periphery of the dorso-longitudinal muscles (DLMs) of the indirect flight musculature (Chaturvedi *et al*. 2017; Boukhatmi and Bray 2018). These are the adult Muscle Stem Cells (MuSCs), also referred to as “satellite cells”, due to their position at the outer edge of the muscle fibre. Here we shall refer to them as adult MuSCs. In mammals, these cells have been characterised as individual resident adult stem cells that lie quiescent under the basal lamina, and are required for muscle homeostasis and repair after injury (Scharner and Zammit 2011; Hung *et al*. 2023). Although first observed in vertebrates in electron microscopy studies as long ago as 1961, adult MuSCs had not been thought to be present in *Drosophila* muscle. Now three papers have established their credentials. As well as occupying a comparable position on the periphery of the adult muscle fibre, lineage tracing has evidenced that they contribute to the flight muscles in both homeostasis and after injury (Chaturvedi *et al*. 2017; Boukhatmi and Bray 2018), and there is conserved function of the RACK1 gene in muscle repair (Catalani *et al*. 2022). These *Drosophila* adult MuSCs express the transcription factor Zfh1, whose product is used as a marker of *Drosophila* MuSCs. In mice, the mammalian orthologue of Zfh1, named Zeb1, is also expressed in their MuSCs (Siles *et al*. 2019).

A subtractive hybridisation screen for genes expressed in mesoderm derivatives (Taylor 2000) identified the *Him* gene. *Him* has a striking expression pattern in the embryo. It is expressed in muscle progenitor cells and rapidly decreases as muscle differentiation gets underway (Liotta *et al*. 2007; Elwell *et al*. 2015). During myogenesis *Him* is expressed in the individual AMPs in the embryo abdominal segments (Liotta *et al*. 2007) and in the wing imaginal disc AMP cell population (Soler and Taylor 2009; this paper). Functionally, *Him* can inhibit muscle differentiation in both phases of myogenesis. The Him protein binds to the conserved transcriptional co-repressor Groucho, via a C-terminal tetrapeptide WRPW, and requires WRPW and *groucho* function to inhibit muscle differentiation (Liotta *et al*. 2007). Moreover, both in tissue culture and *in vivo, Him* can also inhibit the activity of the transcription factor Mef2 (Liotta *et al*. 2007), a key regulator of muscle differentiation (Taylor and Hughes, 2017). The striking expression pattern of *Him* in muscle progenitors in the embryo, and subsequently in the adult progenitors (Soler and Taylor 2009), is suggestive of a function in muscle.

Here, using two new *Him* null alleles, we have uncovered roles for *Him* in both the development and maintenance of adult muscle. *Him* loss-of-function causes both a reduction in the number of wing-disc associated AMPs, as well as an age-associated decline in MuSCs in the adult flies, which has a functional consequence. Furthermore, we show a requirement for *Him* in normal jump muscle development and function.

## Results and Discussion

### *Him* is a new marker of *Drosophila* MuSCs

To visualise *Him* expression, we used a HimGFP “minigene” (Figure 1A). This consists of approximately 3.8kb upstream *Him* sequence, followed by N-terminally eGFP tagged Him coding sequence and the *Him* 3’UTR (Liotta *et al*. 2007). As previously reported (Soler and Taylor 2009), *Him* is expressed in the AMPs associated with the L3 wing imaginal disc (Figure 1B), were it is co-expressed with Mef2. We now show that Him is also expressed in the equivalent cells in the T2 mesothoracic leg disc, i.e. the jump muscle progenitor cells, and is again co-expressed with Mef2 (Figure 1C). Thus, Him is a marker of the adult myoblasts during the larval stage.

**Figure 1.**
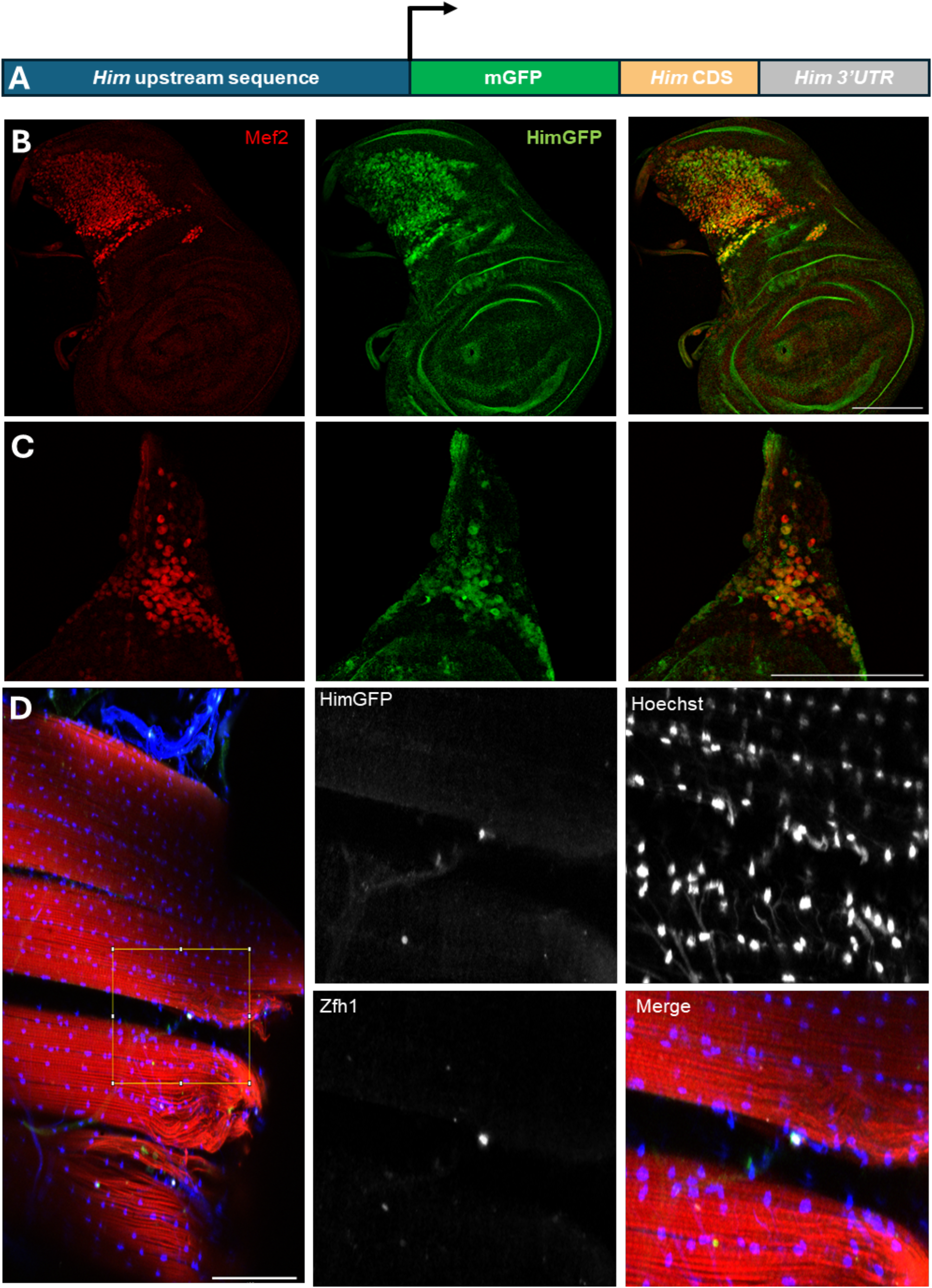
HimGFP design and expression in imaginal disc myoblasts and adult MuSCs. (A) The HimGFP ‘minigene’ consisting of approximately 3.8kb upstream genomic sequence, mGFP fused to the coding sequence of *Him* via an –SSSS-linker, followed by Him’s 3’ UTR. (B) L3 larval wing imaginal disc, labelled with anti-Mef2 and anti-GFP show co-localization of HimGFP and Mef2 in the myoblast population. (C) L3 larval T2 mesothoracic leg discs also show HimGFP and Mef2 co-localization in the associated myoblasts. (D) A sagittal thoracic section showing a close-up of a adult MuSC on the periphery of a dorsal longitudinal muscle fibre, which expresses the canonical marker Zfh1 as well as HimGFP. Scale bars each 100um.

Him expression is known to decline during adult muscle development (Soler and Taylor 2009), but we observed that a small number of isolated cells expressing HimGFP are located on the periphery of the adult DLM fibres. Figure 1D shows that these cells co-express Him and the canonical MuSC marker Zfh1 (Chaturvedi *et al*. 2017; Boukhatmi and Bray 2018). Thus, *Him* is a marker of the novel *Drosophila* adult MuSCs, and only the second gene to show specific expression in these cells.

### Generation of *Him* mutants

We then wished to determine if *Him* has functions in the cells or structures linked to these areas of expression. In order to do this, we made two new mutant alleles of the *Him* gene (Figure 2A). First, *Him*^*0*^ was generated using CRISPR-targeting of the *Him* gene. We isolated an allele with a 2-nucleotide deletion near the N-terminal of the *Him* coding sequence, at codon 25. This produced a frame-shift and results in a premature stop codon at codon 39, and a presumed null allele. Second, *Him*^*52*^ was made by transposable element mediated trans-recombination between two pBac elements, PBac{WH}f06349 and pBac{WH}f04435 (Figure 2B). This produced a small deletion of an approximately 98kb region that includes *Him* plus five other genes. This was verified by PCR analysis: genes between the pBac elements (*Frq1, Him* and *CG33639)* cannot be PCR amplified from *Him*^*52*^ genomic DNA unlike in controls, whereas genes upstream (*Ari1*) and downstream (*Upd2*) of the pBac elements can be amplified from both *Him*^*52*^ and control (Figure 2C). Both the *Him*^*0*^ and *Him*^*52*^ alleles are viable. In our genetic characterisation of the *in vivo* function of *Him*, we also used the previously made HimGFP minigene (Liotta *et al*. 2007) and an available duplication of the *Him* chromosomal region (Dp(1;3)DC343). Thus, we have the tools for an analysis of the role of *Him* in myogenesis.

**Figure 2.**
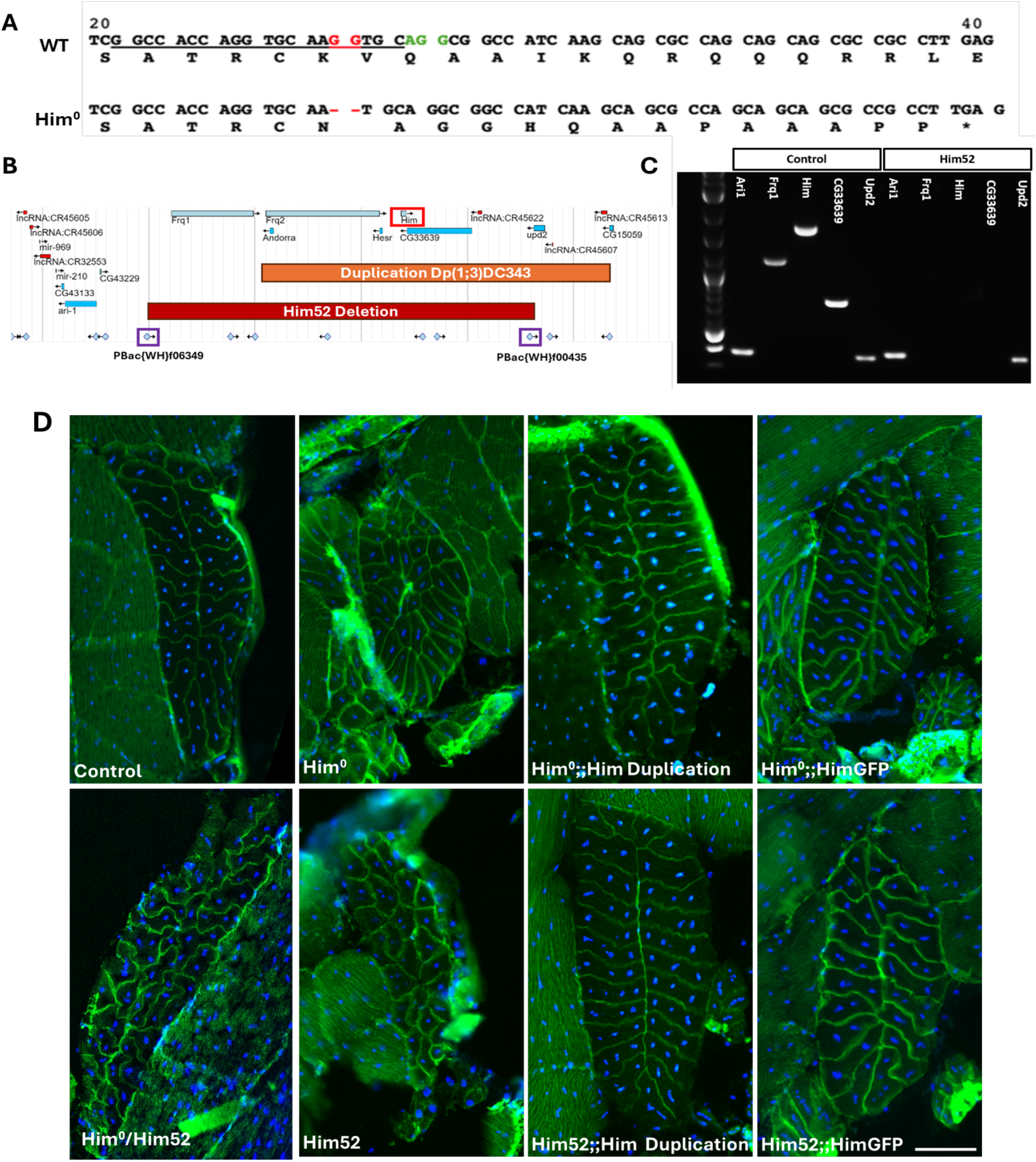
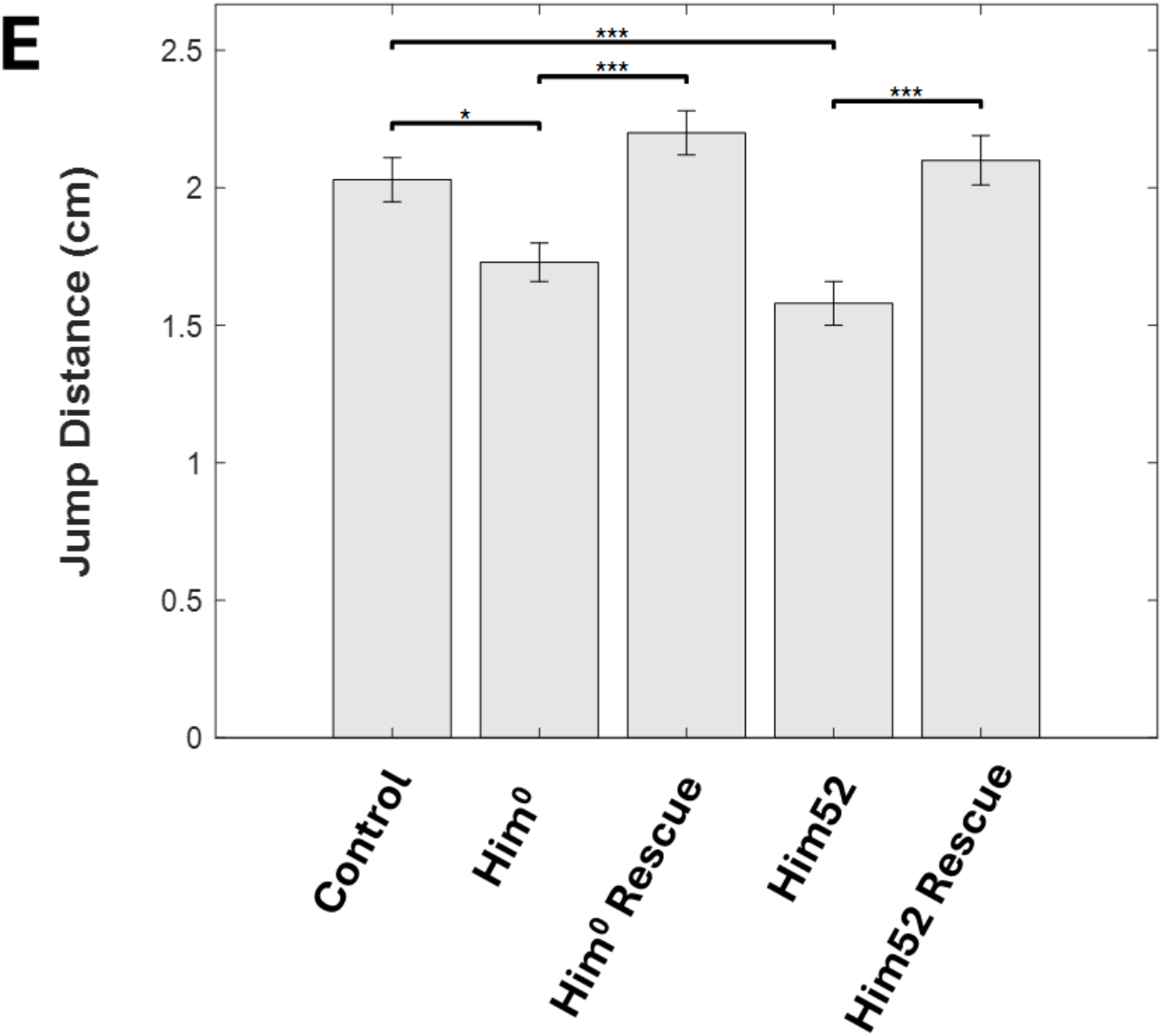
Generation of *Him* mutants and requirement for *Him* in normal jump muscle morphology and function. (A) Partial Wild-type *Him* vs Him ^*0*^ coding sequence before and after CRISPR-mediated 2 nucleotide deletion between codons 25/26. Wild-type sequence shows protospacer (underlined), PAM (green text) and the two nucleotides lost in the *Him*^*0*^ mutant (red). Translations below coding sequences show that the 2-nt deletion causes a premature stop codon. (B) Jbrowse map of the *Him* locus (red box) showing the duplication stock Dp(1;3)DC343 (orange box), which includes 81,651bp DNA spanning *Frq2, Andorra, Hesr, Him, CG33639* and *Upd2*, and the 2 pBacWH FRT elements (purple boxes) used to generate the 98kb *Him*^*52*^ deletion. (C) PCR verification of the *Him*^*52*^ deletion shows amplicons for *Ari1* and *Upd2* outside of the deletion, but not for *Frq1, Him* and *CG33639* within the deletion. (D). Transverse cryosections of young adult jump muscle labelled with integrin betaPS antibody and Hoechst. Wild-type (yw;;nos-Cas9) control jump muscle section consists of two neatly organized rows of muscle fibres. The wild-type morphology is disrupted in 100% of *Him*^*52*^ mutant and *Him*^*0*^ mutant flies, as well as in a heteroallelic mutant combination. One copy of the HimGFP minigene restores wild-type morphology in both *Him*^*52*^ and *Him*^*0*^ mutant background. *Him*^*0*^ and *Him*^*52*^ are both rescued by the *Him* duplication allele. (E) A bar graph showing mean horizontal displacement jumped ± SEM, in response to a looming stimulus. *Him*^*0*^ and *Him*^*52*^ are significantly worse at jumping than yw;;nosCas9 controls. (ANOVA followed by Post-Hoc Tukey HSD test: control vs *Him*^*0*^ *p=0.033, control vs *Him*^*52*^ ***p=0.0003). One copy of HimGFP rescues the *Him* mutant phenotypes to wild-type (ANOVA followed by Post-Hoc Tukey HSD test: *Him*^*0*^ vs *Him*^*0*^;;HimGFP/+ ***p=0.002, *Him*^*52*^ vs *Him*^*52*^;;HimGFP/+ ***p=0.0002, control vs *Him*^*0*^;;HimGFP/+ non-significant, control vs *Him*^*52*^;;HimGFP/+ non-significant).

### *Him* is required for normal jump muscle morphology

The jump muscle is a tubular muscle found in the adult thorax, and comprises two columns of constituent muscle fibres neatly organised in an ovoid pattern around a mid-line. These fibres stretch from the mesothoracic leg (T2) to the dorsal notum of the thorax, and contract to power the force required for the insect to jump (Trimarchi and Schneiderman 1995). The arrangement is stereotypical, although the number of fibres that make up the jump muscle varies depending on the wild-type strain studied (Jaramillo *et al*. 2009).

We found that the *Him* mutant alleles had a jump muscle morphology phenotype: 100% of animals studied showed severe disruption to this ordered pattern. Although the jump muscle still develops, the stereotypical arrangement of the muscle fibres seen in cross-sections is disrupted in both *Him*^*0*^ and *Him*^*52*^ mutants, when compared to control flies. Instead of two neat columns of muscle fibres, the muscle has a fractured, disorganized appearance (Figure 2D). This phenotype is fully penetrant, with all *Him* mutant alleles and allelic combinations displaying a dysmorphic jump muscle structure. Thus, *Him*^*0*^ and *Him*^*52*^ look similar. The heteroallelic combination of the CRISPR allele (*Him*^*0*^) over the *Him* mini-deletion (*Him*^*52*^) has a similar phenotype. Furthermore, both *Him*^*0*^ and *Him*^*52*^ alleles can be rescued to wild type morphology by the HimGFP minigene, and also by the duplication of the *Him* chromosomal region.

Taken together, these results show that the *Him* loss-of-function phenotype is not due to any off-target effect in either mutant allele, that the *Him*^*0*^ allele is a null as intended, and that *Him* loss-of-function can be effectively rescued with the HimGFP transgene. They also show that the HimGFP mini-gene functions like the wild type gene. Thus, we have developed robust reagents for the analysis of *Him* function. Importantly, we used these reagents to establish that the *Him* gene is required in order to make a jump muscle with the normal, ordered arrangement of muscle fibres.

Since Him has been characterised as an inhibitor of Mef2 activity (Liotta *et al*. 2007; Soler and Taylor 2009), these morphological changes observed in the Him mutants may arise from enhanced activity of Mef2. Indeed, Mef2 over-expression also causes defects in jump muscle patterning (Vishal *et al*. 2023).

### *Him* is required for jump muscle function

To determine if the loss of jump muscle organisation has a phenotypic consequence, we undertook a jumping behavioural assay on young adult male flies (<3 days). *Drosophila* jump both as an escape response from a looming stimulus, or to “take-off” to initiate flight (Zumstein *et al*. 2004; Card and Dickinson 2008). Here we used a paintbrush as the looming stimulus and found that both *Him*^*0*^ and *Him*^*52*^ mutant flies had a significant 15-25% reduction in jumping ability compared to control flies (Figure 2E). This demonstrates a functional consequence of the disrupted muscle morphology.

Similarly to the morphological assay, one copy of the HimGFP minigene could rescue the behavioural phenotype of both *Him* alleles to wild-type. The *Him* gene is therefore required for both jump muscle morphology and function.

### *Him* mutants have a reduced adult myoblast pool

We also analysed the wing imaginal discs of control and Him mutant animals, to determine if there were any observable consequences of loss of Him function. While premature muscle differentiation can be a marker for un-restrained Mef2 activity (Lovato *et al*. 2005; Soler and Taylor 2009), we did not observe extensive F-actin accumulation in mutant discs (data not shown). However, we did observe a reduction in the size of the myoblast pool in Him^0^ mutant animals (Figure 3B), and this phenotype was fully rescued by Dp(1;3)DC343 (Figure 3C, and quantified in Figure 3D). This reduction in the myoblast pool was at least partially caused by a reduction in myoblast proliferation, based upon staining for phospho-histone H3 (Figure 3E-H). Given a role for *Him* in inhibiting myoblast differentiation, reduced proliferation in the *Him* mutants might be an indication that the myoblasts are transitioning into a differentiation state.

**Figure 3.**
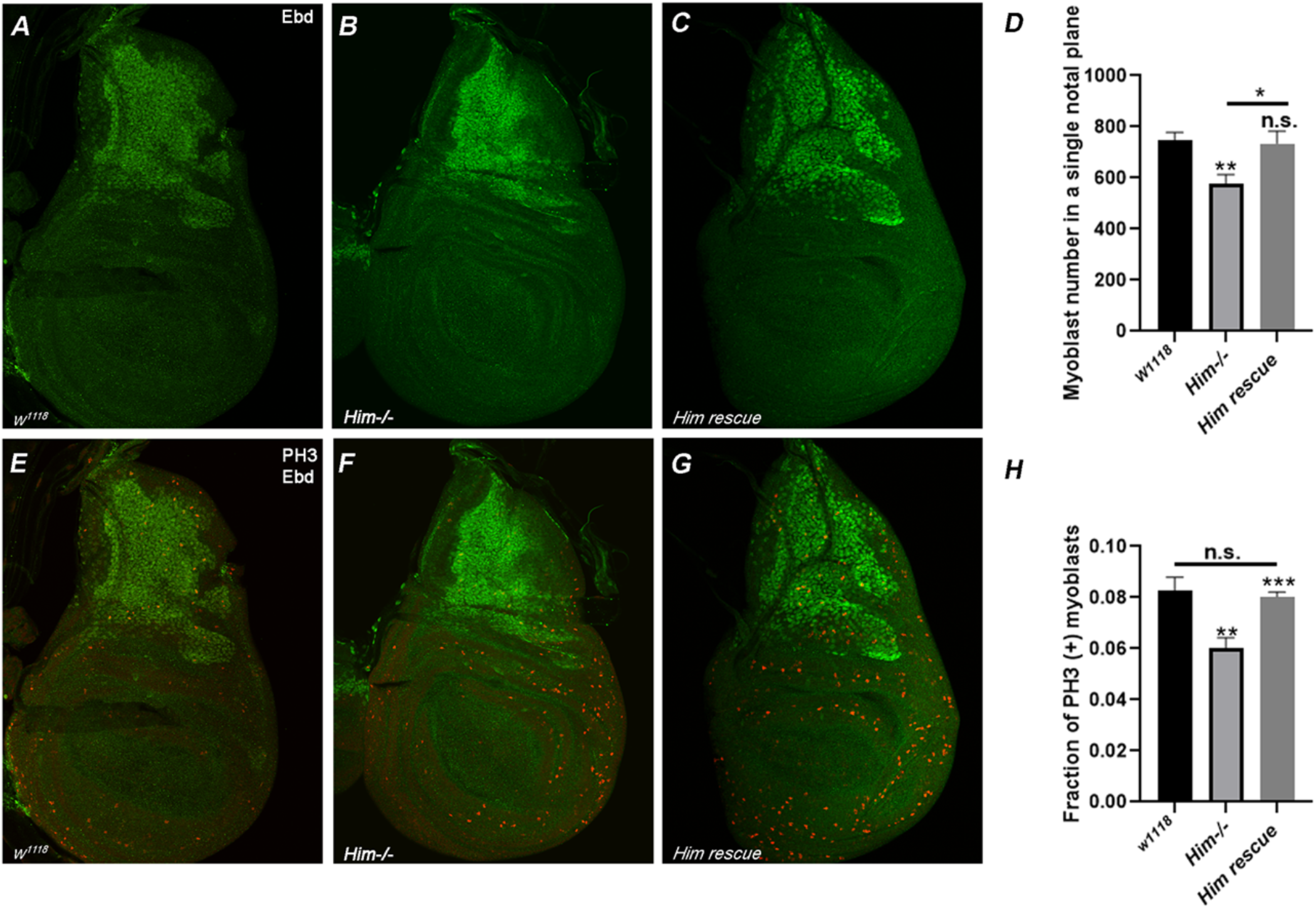
*Him* mutants have a reduced adult myoblast pool. (A-C) Third instar larval wing imaginal discs were stained for Ebd1 (labels all myoblasts) and (E-G) Ebd1 and phospho-histone H3 (PH3), which labels mitotic cells. The myoblast pool size was reduced in *Him*^*0*^ mutants (B) compared to controls (A), and restored when mutants were combined with Dp(1;3)DC343 (C) (ANOVA followed by Post-Hoc Tukey HSD test: Control vs *Him*^*0*^ **p=0.0089, *Him*^*0*^ vs rescue *p=0.0283, control vs rescue non-significant). The fraction of PH3 expressing cells was reduced in *Him*^*0*^ mutants (F) compared to controls (E), and restored when mutants were combined with Dp(1;3)DC343 (G) (ANOVA followed by Post-Hoc Tukey HSD test: Control vs *Him*^*0*^ **p=0.0028, *Him*^*0*^ vs rescue *p=0.0153, control vs rescue non-significant). Myoblast pool size is quantified in panel (D), and fraction of mitotic cells quantified in (H).

### *Him* is required to maintain adult MuSC number and flight ability during ageing

We then asked whether *Him* function affected the number of adult MuSCs, identified with Zfh1 expression in the DLMs. In young flies, the number of MuSCs is similar in *Him*^*0*^ mutants as in controls (Figure 4A). However, although the number of MuSCs in wild type flies was unchanged after ageing to two weeks, strikingly in *Him* mutant flies we found a significant reduction of approximately 50% in Zfh1-positive MuSC number in standard DLM cross-sections (Figure 4A). This was rescued by the duplication to close to the value observed in young *Him* mutant flies.

**Figure 4.**
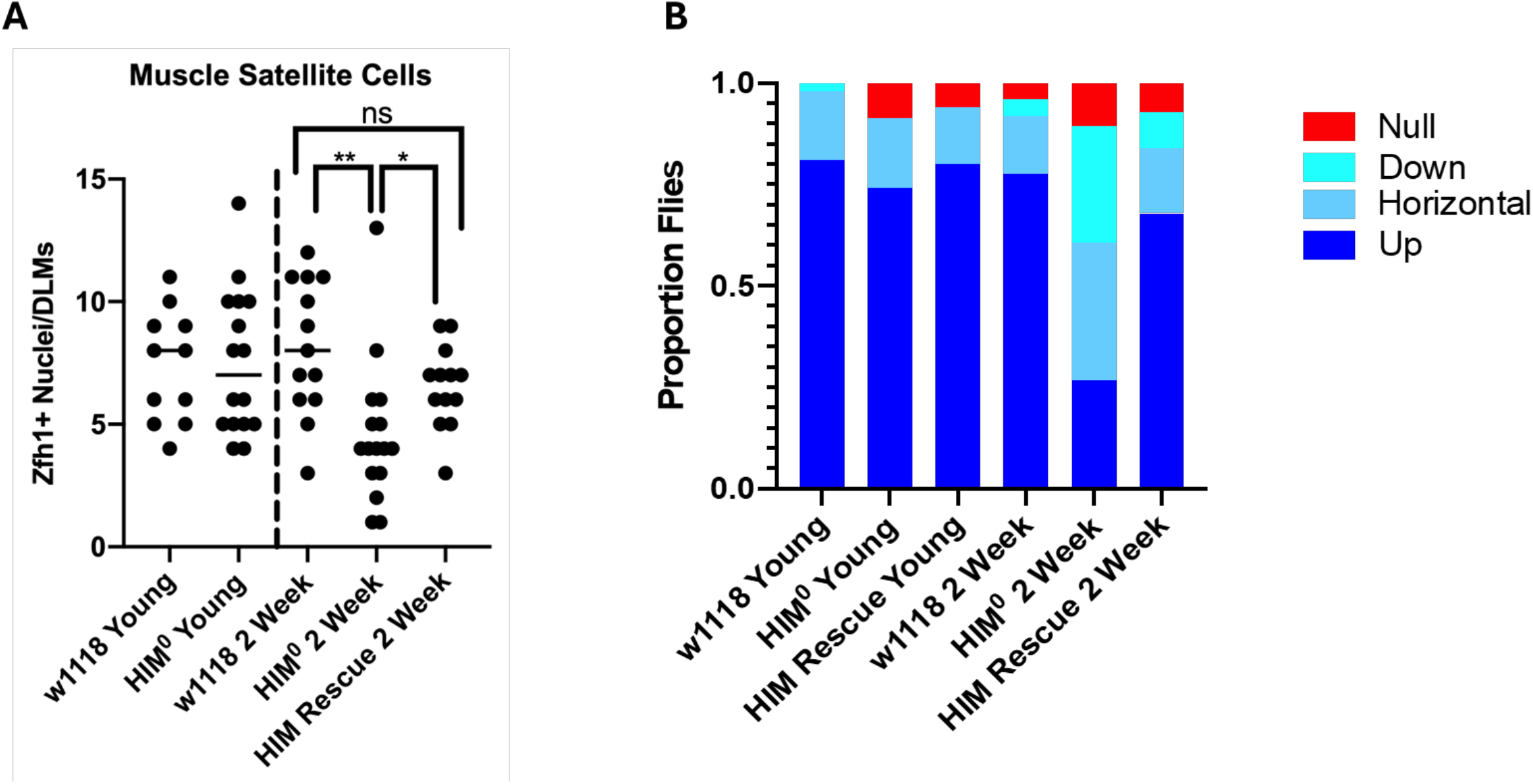
*Him* mutant adult MuSC number and flight ability decrease with age. (A) Flies from control, *Him*^*0*^ mutant, and *Him*^*0*^ mutant combined with duplication, were sectioned and stained to quantify the numbers of adult MuSCs in young and two-week old flies. MuSC number was similar for control and mutants in young flies, however MuSC number was significantly reduced in mutants at two weeks of age (Two-tail t-test: 2-week Control vs 2-week *Him*^*0*^ ** p<0.01, *Him*^*0*^ vs rescue *p<0.05, control vs rescue non-significant). (B) Flies of the same genotypes and ages as in A were flight-tested and scored for whether they flew Upwards, Horizontal, Down, or Not-at-all (Null). Control flies showed strong flight ability as young flies and when aged; and *Him*^*0*^ mutants showed an age-dependent loss of flight ability, which was rescued by Dp(1;3)DC343.

We then asked whether there was any functional consequence to this. Do *Him* mutants show a loss of flight ability over time? We measured flight ability in both young and two-week aged flies using a standard flight assay that scores whether flies fly Upwards, Horizontally, Downwards or Not-at-all, when released below a light in a plexiglass box (Drummond et al. 1990). Control w1118 flies predominantly fly upwards, both as young flies and after ageing for two weeks (>75% in both categories fly upwards; Figure 4B). For the *Him*^*0*^ mutants, young adults showed strong flight ability similar to controls (>70% flew upwards). However, after two weeks only approximately 25% of *Him*^*0*^ adults showed strong flight, and the rest were moderately to severely impaired in their flight ability. The flight-reduced phenotype of the *Him*^*0*^ mutant was rescued by the *Him* duplication, confirming that this effect is due to mutation of the *Him* gene. Thus, *Him* is required to maintain flight ability during ageing.

Taken together, our studies identify multiple roles for the myogenic inhibitor gene *Him* in muscle development and function. *Him* is required to maintain the muscle stem cell pool both at the larval stage and in the adult. We show that in adult *Him* mutants the loss of MuSCs with ageing is accompanied by age-dependent reduction in flight ability. Furthermore, *Him* is required for normal jump muscle organisation and function. One potential mechanism to explain these phenotypes is that Him can inhibit Mef2 activity (Liotta *et al*. 2007). Mef2 is a driver of muscle differentiation (Taylor & Hughes 2017), whereas genes such as *ZD1* and *Him* can inhibit the myogenic program (Postigo *et al*. 1999; Liotta et al 2007). It is striking that whilst Zfh1 levels decline in the majority of wing disc AMPs as development progresses, a subset of cells retain Zfh1 expression and fail to differentiate; these are the MuSCs (Chaturvedi *et al*. 2017; Boukhatmi and Bray, 2018). Like Zfh1, Him is prominently expressed in wing disc-derived myoblasts around the DLM templates at about 20 hours after pupal formation, as the DLMs begin to develop, but disappears by 30 hours (Soler and Taylor 2009; Chaturvedi *et al*. 2017). It is possible that Him functions alongside Zfh1 to repress the myogenic program. As such, loss of *Him* function could cause enhanced Mef2 activity, which in adult MuSCs might push them towards differentiation. This would be consistent with the observations that enhanced Mef2 function in myoblasts phenocopies *Him* loss-of-function in both interfering with jump muscle organization (Vishal *et al*. 2023) and reducing the myoblast pool size (Trujillo *et al*. 2024).

We also note that Him is only the second identified marker, after Zfh1, for *Drosophila* adult MuSCs. As well as making Him a useful tool for the identification of these cells, it provides new knowledge about their biology. Notably, *Him* is the first gene identified that has been demonstrated to be required for maintenance of both adult *Drosophila* MuSCs and flight muscle function. Our studies illustrate the promise of the *Drosophila* system for a genetic dissection of these important cells in the context of the maintenance of developed muscle, as well as of repair following injury, and of muscle deterioration associated with age and disease.

## Methods

### Fly Stocks and Genetics

The following *Drosophila melanogaster* stocks were used: HimGFP (Liotta *et al*. 2007), *Him* duplication Dp(1;3)DC343 (Bloomington Stock Center 32277), yw;;nos-cas9 (BestGene) and w^1118^. *Him*^*0*^ and *Him*^*52*^ were generated during this study.

For rescue experiments, *Him* mutant females were crossed to either HimGFP or *Him* duplication males. The resultant F1 males which were assayed retain a *Him* mutant background with the rescue allele on the third chromosome. Stocks and crosses were maintained at 25°C on standard culture media.

The *Him*^*52*^ line was generated using FRT mediated recombination between the f06349pBacWH and f00435 pBacWH elements, resulting in a 98Kb deletion spanning six genes including *Him*. PCR primer pairs were designed to confirm that the genes between the two pBac elements were deleted, and that genes outside the elements remain intact. The genes analysed and their respective primer pairs included *Ari1* (forward 5’-GCCTGGAGCCACCTTCTAG, reverse 5’-CAGTCACAAGGCTCAGAAGC), *Frq1* (forward 5’-CTTTCCACAAGGCGATCCCA, reverse 5’-CAGGTTGCCCTTCGATGTG), *Him* (forward 5’-AGTTTGGCGCGCAATGTG, reverse 5’-GAACGAAGGCAGATGGAG), *CG33639* (forward 5’-GGATCTACGTTCATTCATTGTG, reverse 5’-CGAGGACTAAGTTATCTAGTTACAT) and *Upd2* (forward 5’-CTCGCTATAATACTGGAAGGGC and reverse 5’-GAGCGTCTCAACTTTCACACG).

The Him^0^ line was generated using CRISPR/Cas9 genome editing. Flies of the genotype yw; nos-Cas9 were injected with a sgRNA (IDT) targeting Him (protospacer sequence underlined in Figure 2A). G0 adults arising from the injected embryos were separately crossed to generate stable lines, and each line was screened for mutations in Him using PCR followed by sequencing. One line, designated Him?, had a 2-nt deletion beginning in codon 25, resulting in a frameshift and subsequent premature stop codon at position 39.

### Immunofluorescence and confocal microscopy

Wing and leg imaginal discs were dissected from wandering third instar larvae, fixed and immuno-stained as described by Klipa *et al*. 2023. Jump muscle cryo-sections were obtained from young adult flies (<2 days) as described by Morriss *et al*. 2012.

To image adult MuSCs, sagittal thoracic DLM sections were prepared as previously described (Leroux *et al*. 2023) with minor modification. Adult flies (<2 days) were anesthetized using CO_2_ and had their heads, abdomens, legs and wings removed, before fixation in 4% PFA/1% Triton in PBS for 20 minutes at room temperature. Thoraces were bisected with a scalpel, followed by a further fixation for 25 minutes at room temperature. Samples were washed 3×20 minutes with 0.3% PBT, and then blocked in 0.5% BSA/0.3% PBT for half-hour. Primary antibodies were diluted in blocking solution, and incubated with the sample overnight at 4°C with agitation. The following morning, samples were washed 3×20 minutes with 0.3% PBT and then incubated with secondary antibodies diluted in blocking solution for 2 hours at room temperature with agitation. Secondary antibodies were removed, and samples were washed for 3×20 minutes with PBS prior to mounting on glass slides in Vectashield.

Antibodies used for immunolabelling include guinea pig anti-Ebd 1:2000 (Yashi Ahmed), rabbit-anti-Mef2 1:1,000 (DSHB), rabbit-anti-Zfh1 1:5,000 (Ruth Lehman), chicken-anti-GFP 1:500 (Abcam, ab13970) and mouse anti-βps-integrin 1:50 (DSHB). Secondary antibodies were anti-chicken-488 (Jackson Immuno, 703-545-155), anti-rabbit-555 (Thermofisher, A-21428) and anti-mouse-488 (Thermofisher, A-11029). Nuclei were stained using DAPI 1:1,000 (Fisher, 62248), and mitotic nuclei labelled with PH3 (1:100, Millipore). Samples were imaged on an Olympus BX63 fluorescent microscope, or a LSM880 confocal microscope.

### Functional Jumping and Flight assays

Jumping ability was tested according to Zumstein *et al*. 2004 with minor modification. Young (<2 days) male flies were anaesthetized on a fly pad using CO2, and had their wings removed using dissection scissors. Flies were returned to a culture vial at 25°C overnight, to recover from being anesthetized. Individual flies were transferred onto a piece of lined paper for a jumping trial, which was filmed using a smart phone. A paintbrush was used as a looming stimulus to elicit a jump escape response, by moving the brush towards the posterior of the fly. The horizontal distance travelled by the jumping fly was calculated using Tracker software (physlets.org/tracker). Each fly had 5 jumps recorded, with the mean of the longest 3 calculated and used for comparison. A minimum of 15 flies per genotype were assayed. Flight testing was performed as according to Drummond *et al*. 1991 using at least 50 flies per genotype.

### Quantification of adult muscle stem cells

Wing disc associated muscle stem cells were quantified as described in Vishal *et al*. 2023. DLM-associated MuSC numbers were quantified using transverse thoracic cryo-sections. MuSCs were identified on the periphery of muscle fibres using Zfh1 antibody staining. At least ten animas were analysed for each genotype and time point.

## Notes

### Competing Interest Statement

The authors have declared no competing interest.

## References

Boukhatmi, H. and Bray, S. (2018). A population of adult satellite-like cells in Drosophila is maintained through a switch in RNA-isoforms. eLife 7:1–24. doi: 10.7554/eLife.35954.

Card, G. and Dickinson, M.H. (2008). Visually mediated motor planning in the escape response of Drosophila. Current biology : CB 18(17):1300–1307. doi: 10.1016/J.CUB.2008.07.094.

Catalani, E., Zecchini, S., Giovarelli, M., Cherubini, A., Del Quondam, S., Brunetti, K., … Cervia, D. (2022). RACK1 is evolutionary conserved in satellite stem cell activation and adult skeletal muscle regeneration. Cell Death Discovery 2022 8:1 8(1):1–14. doi: 10.1038/s41420-022-01250-8.

Chaturvedi, D., Reichert, H., Gunage, R.D. and VijayRaghavan, K. (2017). Identification and functional characterization of muscle satellite cells in Drosophila. eLife 6. doi: 10.7554/ELIFE.30107.

Dobi, K.C., Schulman, V.K. and Baylies, M.K. (2015). Specification of the somatic musculature in Drosophila †. doi: 10.1002/wdev.182.

Drummond, D.R., Hennessey, E.S. and Sparrow, J.C. (1991). Characterisation of missense mutations in the Act88F gene of Drosophila melanogaster. MGG Molecular & General Genetics 226(1–2):70–80. doi: 10.1007/BF00273589.

Elwell, J.A., Lovato, T.A.L., Adams, M.M., Baca, E.M., Lee, T. and Cripps, R.M. (2015). The myogenic repressor gene Holes in muscles is a direct transcriptional target of Twist and Tinman in the Drosophila embryonic mesoderm. Developmental Biology 400(2):266–276. doi: 10.1016/j.ydbio.2015.02.005.

Hung, M., Lo, H.F., Jones, G.E.L. and Krauss, R.S. (2023). The muscle stem cell niche at a glance. Journal of Cell Science 136(24). doi: 10.1242/JCS.261200/337756.

Jaramillo, M.A.S., Lovato, C. V., Baca, E.M. and Cripps, R.M. (2009). Crossveinless and the TGFβ pathway regulate fiber number in the Drosophila adult jump muscle. Development 136(7):1105– 1113. doi: 10.1242/DEV.031567.

Klipa, O., Gammal, M.El and Hamaratoglu, F. (2023). Elimination of aberrantly specified cell clones is independent of interfacial Myosin II accumulation. Journal of Cell Science 136(13). doi: 10.1242/JCS.259935/316700/AM/ELIMINATION-OF-ABERRANTLY-SPECIFIED-CELL-CLONES-IS.

Laurichesse, Q. and Soler, C. (2020). Muscle development : a view from adult myogenesis in Drosophila. Seminars in cell & developmental biology 104:39–50. doi: 10.1016/J.SEMCDB.2020.02.009.

Leroux, E., Ammar, N., Moinet, S., Pecot, T. and Boukhatmi, H. (2023). Studying Muscle Transcriptional Dynamics at Singlemolecule Scales in Drosophila. Journal of Visualized Experiments 2023(199). doi: 10.3791/65713.

Liotta, D., Han, J., Elgar, S., Garvey, C., Han, Z. and Taylor, M. V. (2007). The Him Gene Reveals a Balance of Inputs Controlling Muscle Differentiation in Drosophila. Current Biology 17(16):1409– 1413. doi: 10.1016/j.cub.2007.07.039.

Lovato, T.L., Benjamin, A.R. and Cripps, R.M. (2005). Transcription of Myocyte enhancer factor-2 in adult Drosophila myoblasts is induced by the steroid hormone ecdysone. Developmental Biology 288(2):612–621. doi: 10.1016/j.ydbio.2005.09.007.

Morriss, G.R., Bryantsev, A.L., Chechenova, M., Labeau, E.M., Lovato, T.L., Ryan, K.M. and Cripps, R.M. (2012). Analysis of Skeletal Muscle Development in Drosophila. Methods in molecular biology (Clifton, N.J.) 798:127. doi: 10.1007/978-1-61779-343-1_8.

Postigo, A.A., Ward, E., Skeath, J.B. and Dean, D.C. (1999). zfh-1, the Drosophila Homologue of ZEB, Is a Transcriptional Repressor That Regulates Somatic Myogenesis. Molecular and Cellular Biology 19(10):7255. doi: 10.1128/MCB.19.10.7255.

Scharner, J. and Zammit, P.S. The muscle satellite cell at 50: the formative years. Skeletal Muscle 1, 28 (2011). 10.1186/2044-5040-1-28

Siles, L., Ninfali, C., Cortés, M., Darling, D.S. and Postigo, A. (2019). ZEB1 protects skeletal muscle from damage and is required for its regeneration. Nature Communications 2019 10:1 10(1):1–18. doi: 10.1038/s41467-019-08983-8.

Soler, C. and Taylor, M. V. (2009). The Him gene inhibits the development of Drosophila flight muscles during metamorphosis. Mechanisms of Development 126(7):595–603. doi: 10.1016/j.mod.2009.03.003.

Taylor, M. V. (2000). A novel Drosophila, mef2-regulated muscle gene isolated in a subtractive hybridisation-based molecular screen using small amounts of zygotic mutant RNA. Developmental Biology 220(1):37–52. doi: 10.1006/DBIO.2000.9608.

Trimarchi, J.R. and Schneiderman, A.M. (1995). Initiation of flight in the unrestrained fly, Drosophila melanogaster. Journal of Zoology 235(2):211–222. doi: 10.1111/J.1469-7998.1995.TB05138.X.

Trujillo, E.M., Lee, S.R., Aguayo, A., Torosian, T.C. and Cripps, R.M. (2024). Enhanced expression of the myogenic factor Myocyte enhancer factor-2 in imaginal disc myoblasts activates a partial, but incomplete, muscle development program. Developmental biology 516:82–95. doi: 10.1016/J.YDBIO.2024.08.004.

Vishal, K., Barajas Alonso, E., DeAguero, A.A., Waters, J.A., Chechenova, M.B. and Cripps, R.M. (2023). Phosphorylation of the Myogenic Factor Myocyte Enhancer Factor-2 Impacts Myogenesis In Vivo. Molecular and cellular biology 43(6):241–253. doi: 10.1080/10985549.2023.2198167.

Zumstein, N., Forman, O., Nongthomba, U., Sparrow, J.C. and Elliott, C.J.H. (2004). Distance and force production during jumping in wild-type and mutant Drosophila melanogaster. Journal of Experimental Biology 207(20):3515–3522. doi: 10.1242/JEB.01181.

